# Set202 is a histone lysine methyltransferase that enables *Cryptococcus neoformans* to adapt to the host environment

**DOI:** 10.64898/2026.09.15.751726

**Authors:** Georgina Agyei, Kayla C. Anderson, Peter V. Stuckey, Federico J. Celis, Timothy J. Theisen, Felipe H. Santiago-Tirado

## Abstract

All pathogens rely on dynamic gene expression for host adaptation and disease causation. This is especially important for pathogens that normally are found in the environment, like *Cryptococcus neoformans,* given that environmental and host conditions are dramatically different. Epigenetic modifications, like those carried out by SET-domain histone lysine methyltransferases (HKMTs), are essential for this genome expression flexibility. While transcription factors and signaling cascades have been well-studied in *C. neoformans*’s host adaptation, the role of HKMTs in this process has not been explored. Here, we characterize the gene *SET202*, previously reported as a putative HKMT important for growth in host lungs. We show that *set202Δ* mutants have 50% less phagosomal permeabilization than the wild-type fungi, have phagosomal maturation defects, and poor intracellular survival. Additionally, they are defective in traits promoting virulence, including significantly smaller capsules, less melanin, and poor growth under host-like conditions. These phenotypes contributed to reduced virulence in a murine model of infection but, notably, this defect was host-temperature-dependent. In order to explain mechanistically how *SET202* deletion could affect such a wide range of processes, we assessed and demonstrated, for the first time, that Set202 is a true histone-3-lysine 36 methyltransferase, supporting its role in epigenetically modifying the fungal genome to enable adaptation to the hostile host conditions. Consequently, our findings support the study of the biological function of Set202, as it can serve as a potential future therapeutic target for the treatment of infections caused by *C. neoformans,* and possibly other pathogenic fungi.

**IMPORTANCE:** Epigenetic modifications are essential for a wide variety of processes, from cancer to infectious diseases. In the fungal world, they have been associated with fungal stress response, antifungal resistance, and disease establishment. However, in the environmental opportunistic pathogen *Cryptococcus neoformans*, the role of histone methyltransferases (HKMTs), one of the main epigenetic regulators, has been poorly elucidated. Herein, we characterize the role of a putative HKMT, Set202, in *C. neoformans’* transition from the environment to the host. This is utmost important because this fungus’s main driver of disease is its ability to adapt to host conditions. We show that Set202 regulates a variety of processes needed for disease by acting as a true HKMT. Given the inadequate treatments available for diseases caused by this fungus, and that HKMTs are conserved in other fungal pathogens, characterizing Set202 function will provide crucial information to develop new treatments for this and other fungal diseases.

## INTRODUCTION

Invasive mycoses are a leading cause of infectious disease-related mortality among the immunocompromised (1, 2). Worldwide, they cause 6.5 million cases annually, with more than half ending in death (3, 4). Central to the pathogenesis of these fungal diseases are the pathogens’ mechanisms to adapt to host conditions (5). Mechanisms for nutrient acquisition, high carbon dioxide concentration resistance, immune evasion, and thermotolerance are indispensable for establishing a niche in a mammalian host, which subsequently leads to disease (6–8). For example, within the *Cryptococcus* genus, thermo-intolerance prevents *Cryptococcus amylolentus* from causing disease in a mammalian host, unlike its close relative, *Cryptococcus neoformans* (9).

*C. neoformans* is an environmental fungus with a worldwide distribution, especially associated with bird guano, soil, and decaying trees (10, 11). During infection, desiccated yeasts or spores are inhaled into the respiratory tract and travel down to the alveoli, where they encounter the first host defenses. In immunocompetent individuals, these inhaled infectious particles become latent or are cleared by host immune cells. However, in immunocompromised individuals, *C. neoformans* disseminates from the lungs and causes cryptococcal meningoencephalitis (CME), an infection of the brain and meninges that is highly lethal (12). This is possible because it undergoes molecular and cellular changes that enable it to survive inside the host (13, 14), including adapting to the host’s higher temperature, higher carbon dioxide concentration, and restricted nutrients, as well as producing virulence factors that allow it to evade the host’s immune defenses (6, 15). Several transcription factors and signaling cascades have been implicated in these adaptation processes, but the role epigenetic regulators play in rapidly modulating gene expression during the transition from the environment to the host is less clear. Histone lysine methyltransferases (HKMTs) are a major group of epigenetic regulators that belong to the Su(var)3-9, Enhancer of Zeste, Tri-thorax (SET) domain-containing proteins (16). Named after their orthologs in *Drosophila melanogaster*, the SET domain proteins are conserved across Eukarya (16, 17). In *Saccharomyces cerevisiae*, a non-pathogenic fungus, these proteins are crucial for maintaining transcriptional fidelity under nutrient stress (18). Among pathogenic fungi, they have been associated with fungal stress responses, antifungal resistance, disease dissemination, and virulence (19–22).

A previous screen aimed at understanding the pathogenesis of *C. neoformans* found that deletion of the gene *SET202* in the H99 clinical strain resulted in poor lung survival in murine infections (23). The Set202 protein was also shown to be essential for melanin production, an important cryptococcal virulence factor that protects the fungus from reactive oxygen species-mediated damage in immune cells. Furthermore, previous work from our lab showed that *SET202* is involved in fungal evasion of phagocytosis (24). However, despite these findings, the actual cellular function of Set202 has not been elucidated.

Here, we investigated the role of *SET202* in fungal adaptation to host conditions and its contributions to cryptococcal pathogenesis. We found that *SET202* deletion significantly delayed disease progression relative to wild-type (WT) in our mouse inhalational disease model. We believe this is a consequence of defective host adaptation resulting in slowed growth, altered virulence factor production, and impaired calcium homeostasis. Moreover, *set202Δ* mutants were less able to survive intracellularly in macrophages, probably due to an inability to prevent phagosomal maturation. Lastly, we demonstrate, for the first time, that Set202 is a true H3K36MT, providing a mechanistic basis for these phenotypes. Together, our data strongly support the biological function of *SET202* in epigenetically regulating the fungal transition from the environment to the host, driving disease establishment and progression, and providing insights into possible future therapeutic targets for the treatment of CME.

## RESULTS

### Deletion of *SET202* impairs phagosomal membrane damage

In a prior study, we determined that over 20% of the *C. neoformans*-containing phagosomes (CCP) are actively damaged by the fungus in THP-1 immune cells, a phenomenon important for fungal virulence that has also been reported in murine BMDM (25, 26). Adopting this assay, we used an anti-galectin-3 antibody as a proxy for measuring CCP membrane damage (**Fig. 1A**) and screened for fungal genes involved in this virulence process. We examined 50 fungal mutants known to exhibit altered phagocytosis by THP-1 cells (24), scoring those CCPs with no membrane damage **(Fig. 1B**, top) and those with damage (**Fig. 1B**, bottom), which have a halo of anti-galectin-3 enrichment around the fungus. In general, we found that high uptake mutants have significantly less CCP membrane damage compared to the WT KN99, but more than the nonpathogenic yeast control, *Saccharomyces cerevisiae* (**Fig. 1C, Table S1**). Notably, we found the reverse to be true for the low uptake mutants, which caused CCP membrane damage similar to that of WT (**Fig. 1D, Table S1**).

**Fig 1.**
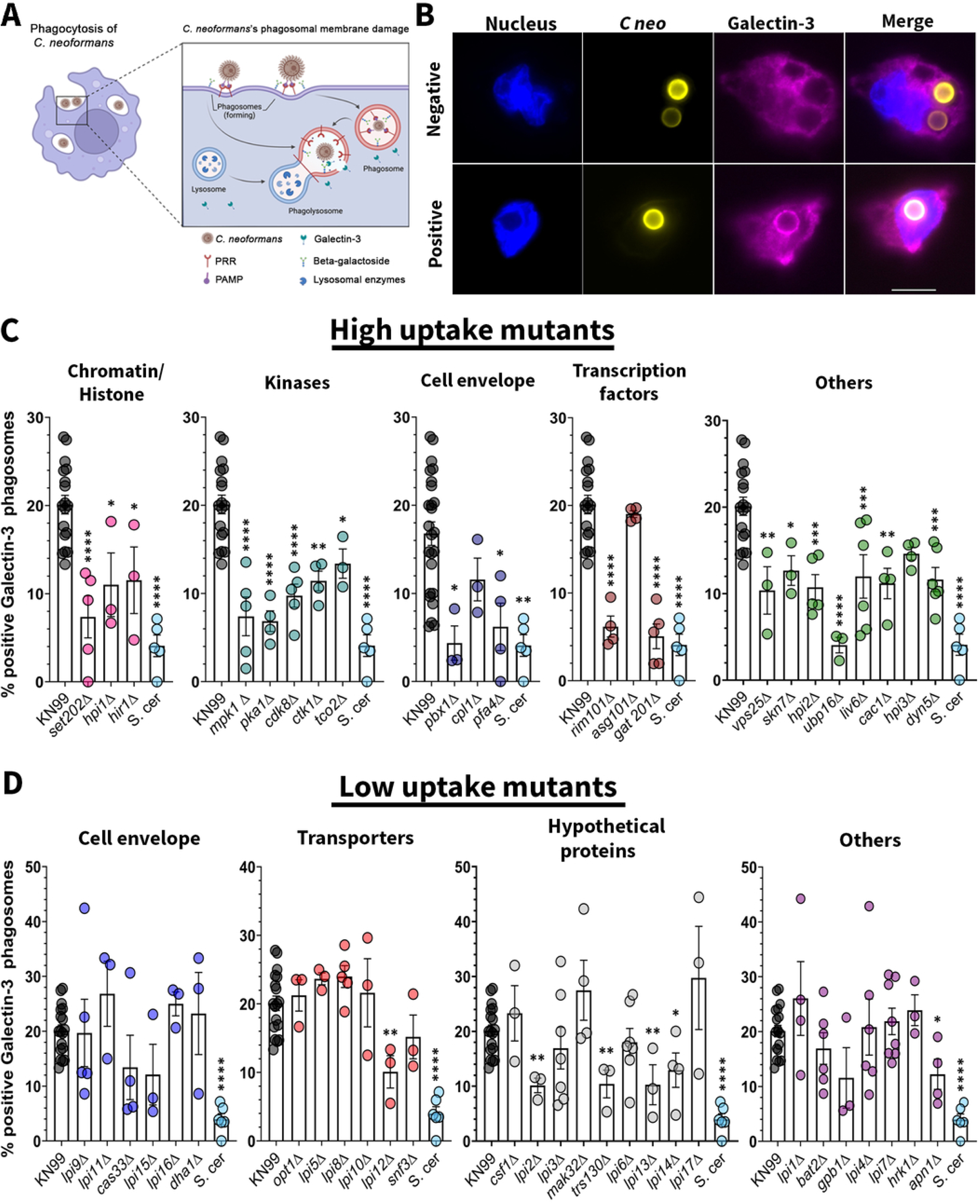
*set202Δ* mutants fail to induce phagosomal membrane damage. (A) Illustration describing the basis of using galectin-3 staining as a proxy for cryptococcal phagosomal membrane damage. This assay was used to screen a small collection of mutants for genes needed for phagosomal permeabilization. PRR, pattern-recognition receptor; PAMP, pathogen associated molecular pattern. (B) Representative images for negative and positive galectin-3 staining depicting membrane damage in infected THP-1 cells. Nucleus of THP-1 cells, blue; *C. neoformans*, yellow; galectin-3, purple; scale bar, 10 microns. (C and D) Quantification of cryptococcal phagosomes positive for galectin-3 staining after 8 hours in naive THP-1 macrophages infected with WT (KN99, first black bar in C and D) and indicated mutants. (C) High-uptake mutants including *set202Δ* (pink). (D) Low uptake mutants, compared to *S. cerevisiae* (last cyan bar in C and D). Significance was determined using one-way ANOVA, comparisons made to KN99; *, P<0.05; **, P<0.01; ***, P<0.001; ****, P<0.0001. Panel A created in BioRender (https://biorender.com/7gkcxm6).

Since CCP membrane damage is associated with virulence, mutants of genes that showed less of this damage possibly play important roles in this process. We therefore decided to take a closer look at the high uptake mutant screen. Most of these have been well characterized in the field, with several of them involved in important fungal virulence processes. However, we found that there is very little data on the chromatin/histone subgroup mutants.

Several recent studies have shown that chromatin epigenetic modifications are important for virulence mechanisms like fungal adaptation, antifungal drug resistance, and disease dissemination (19, 22); thus, we decided to focus on the gene *SET202* that encodes a putative histone methyltransferase that has previously been implicated in virulence, yet with an unknown cellular function (23, 27).

### *SET202* delays CCP maturation, contributing to intracellular fungal survival

Given that *SET202* is essential for CCP membrane damage, we next assessed if it also played a role in the phagosomal maturation process by quantifying colocalization of the vacuolar ATPase (vATPase; **Fig. 2A**) and lysosome-associated membrane protein-1 (LAMP-1; **Fig. 2B**) with the CCP. We found that phagosomes containing *set202Δ* mutants have a higher recruitment of v-ATPase at 30 minutes and 1 hour compared to the WT and complemented strains (**Fig. 2C**). However, we found that there was no difference in the recruitment of LAMP-1 to phagosomes containing *set202Δ* mutants, WT, or complemented strains at 2 hours and 4 hours **(Fig. 2D**).

**Fig 2.**
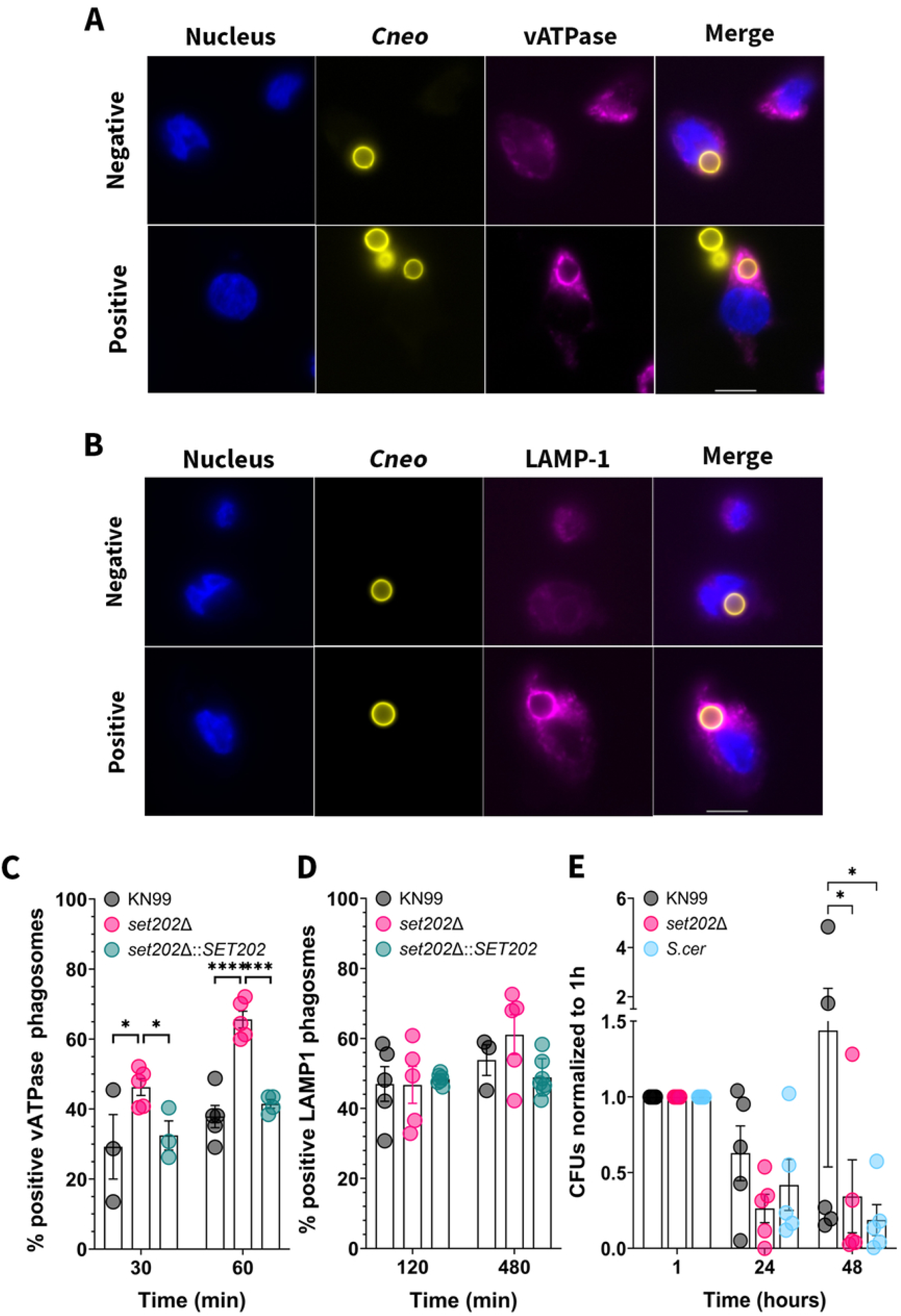
*set202Δ* mutants have low intracellular survival in host immune cells. (A and B) Representative images for negative (top) and positive (bottom) examples of v-ATPase (A) or LAMP-1 (B) colocalization with the cryptococcal-containing phagosome in naïve THP-1 cells. Nucleus of THP-1 cells, blue; *C. neoformans*, yellow; vATPase/LAMP-1, magenta. Scale bar, 10 microns. (C and D) Quantification of cryptococcal phagosomes positive for vATPase (C) or for LAMP-1 (D) among indicated strains. (E) Fungal killing assay showing CFUs of the indicated strains over 48 hours normalized to 1 hour. Significance was determined using two-way ANOVA with comparisons made to *set202*Δ (C and D) or to KN99 (E). *, P<0.05; ****, P<0.0001.

Next, we tested for intracellular survival in THP-1 macrophages. We found that set*202Δ* mutants are readily killed by THP-1 macrophages and behave similarly to nonpathogenic *S. cerevisiae* (**Fig. 2E**). Collectively, these data suggest that *SET202* is essential for fungal immune evasion in the host.

### *SET202* mediates normal disease progression in murine models

So far, we have established that *SET202* plays an important role in intracellular fungal survival by altering phagosomal membrane damage and maturation (**Figs. 1 and 2**). We reasoned that these defects may affect disease progression *in vivo*; thus, we tested the virulence of the mutant in an intranasal murine model of infection. Mice infected with *set202Δ* mutants had prolonged survival, with a median time of death (TOD) of 42.5 days, approximately 2.5 times longer than the 18-day median survival seen in WT and complement-infected mice (**Fig. 3A**).

**Fig 3.**
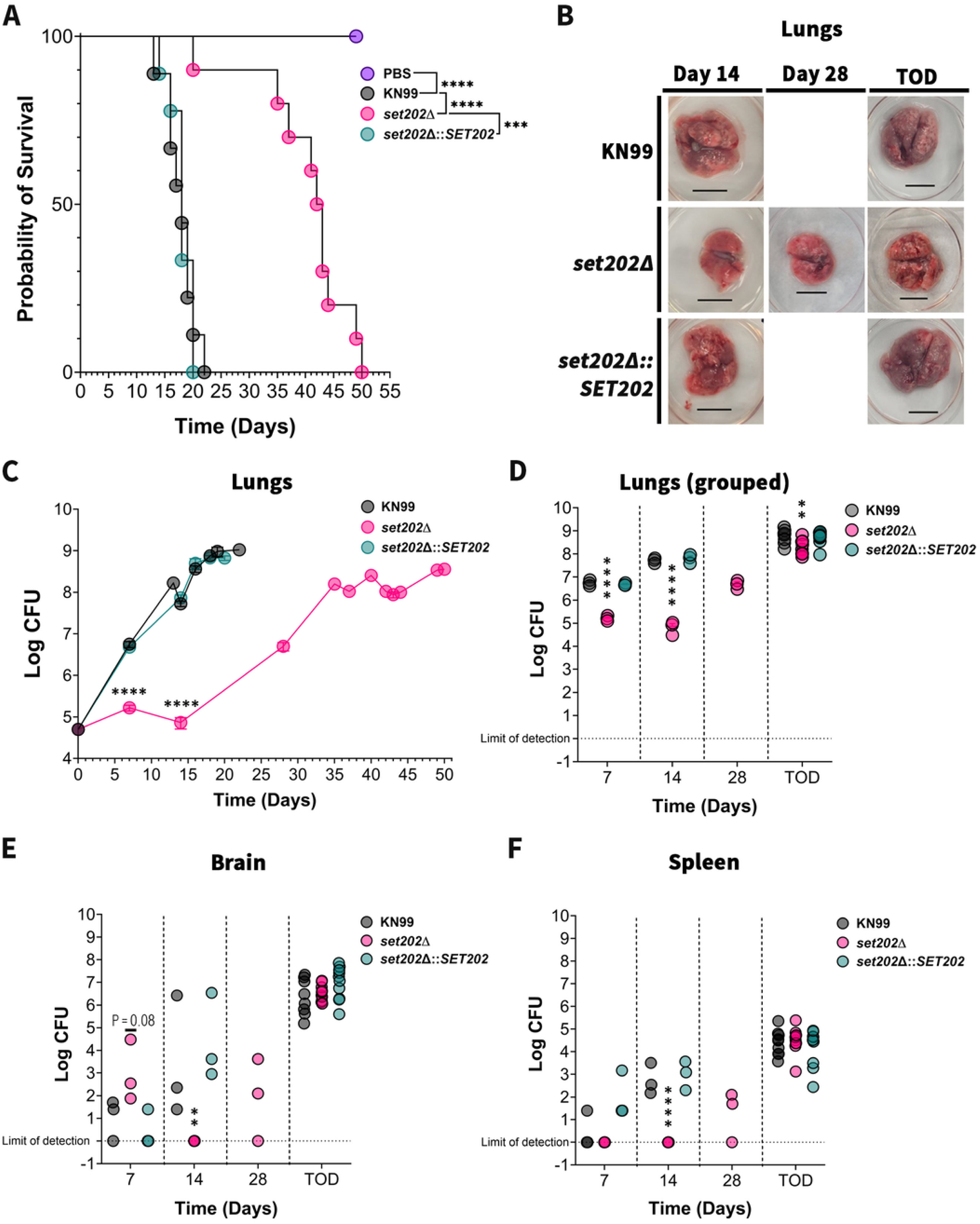
SET202 promotes normal disease progression. (A) Survival curve of A/J mice infected with the indicated strains (N = 4 to 10 per group; inoculum: 5 × 10^4^ fungal cells). (B) Dissected mouse lungs from indicated strains at days 14, 28, or at time of death (TOD). Scale bar,10 mm. (C – F) Organ fungal burden quantified over time in lungs (individual timelines in C and TOD combined in D), brain (E), and spleen (F). Significance was determined with the Mantel-Cox test (A) or the Log-normal one-way ANOVA compared to *set202*Δ (C – F). **, P<0.01; ****, P<0.0001.

Given this increased survival and the previously reported defect of *set202Δ* mutants in murine lungs (23), we assessed the fungal burden in the lungs, brain, and spleen over time (**Figs. 3B-F**). In the lungs, *set202Δ* mutants had an initial prolonged fungistatic phase, with burden at days 7 and 14 staying around ∼10^5^ CFUs, about 2-3 logs less than the WT and complemented strains (**Fig. 3C and D**). At day 7, the fungal counts for the *set202Δ* mutants were only about twice the initial inoculum, and only reached day 7 levels of WT and complemented strains at day 28 (**Fig. 3C and D**), supporting the prior observation that these mutants grow poorly in murine lungs (23). Interestingly, we found no difference in lung fungal burden between the three strains at their time of death (TOD), indicating that eventually the *set202Δ* mutants reached lethal fungal loads.

Gross examination of the lungs at day 7 showed healthy lung anatomy for all the strains (data not shown). However, at day 14, the wild-type and complemented strains showed massive surface gross consolidation and nodules (**Fig. 3B**); the *set202Δ* mutant’s lungs, on the other hand, looked healthy, just like at day 7. The *set202Δ* mutant’s lungs only started showing these surface anatomical distortions and consolidation at day 28. This 3-week lag in the lung disease course correlated with the course seen for the mice’s survival (**Fig. 3A**).

In line with the lung findings, *set202Δ* mutants showed corresponding dissemination patterns in the spleen (**Fig. 3F**). No *set202Δ* mutants’ fungal colonies were detected at days 7 and 14. At day 28, *set202Δ* mutants showed similar spleen dissemination patterns as days 7 and 14 for the WT and complemented strains, before they reached the critical lethal fungal counts at TOD, which corresponded with when critical fungal populations were reached in the lungs.

Interestingly, there was an initial sublethal brain dissemination of *set202Δ* mutants at day 7 (**Fig. 3E**). This was cleared by day 14, followed by gradual re-seeding of the brain on day 28, and the final lethal fungal counts at TOD.

Collectively, these results support our prior *in vitro* immune cell findings and show that *SET202* is an important component of the fungal host adaptation machinery, needed for normal disease progression.

### *SET202* mediates fungal growth in host-like conditions

In our *in vivo* studies above, *set202*Δ mutants grew poorly in murine lungs. We reasoned that deleting *SET202* made the fungus less resilient in the host lung environment during transition from the outside world. Several host defensive factors within the lungs protect against pathogenic invasion. These include a higher carbon dioxide concentration (∼5%), higher temperature (37°C), limited nutrients, immune cells, and other antimicrobial factors. To test if the slow growth was due to defects under these *in vivo* conditions, we compared the *set202*Δ mutant’s growth under optimal conditions (nutrient-rich, ambient temperature, and ambient carbon dioxide concentration), which simulated the external environment, with growth in simulated host-like conditions (nutrient-limited/higher temperature/higher carbon dioxide concentrations). We found that on YPD agar plates, incubated at 30°C or 37°C with or without 5% CO_2,_ there were no significant differences among the strains (**Fig. 4A**). Similarly, there were small, albeit noticeable, differences in YNB agar plates at both 30°C and 37°C which became more noticeable with addition of CO_2_ (**Fig. 4B**). However, in liquid YPD media, we found some significant defects in the growth of the *set202Δ* mutant (**Fig. 4C**), which was even more pronounced in host-like RPMI liquid media at 37 °C and 5% CO_2_ (**Fig. 4D and E**). Consistently, we found significant growth defects with other host-like liquid media (**Fig. S1**). Thus, our *in vitro* findings support the slow *in vivo* growth we see with *set202Δ* mutants.

**Fig 4.**
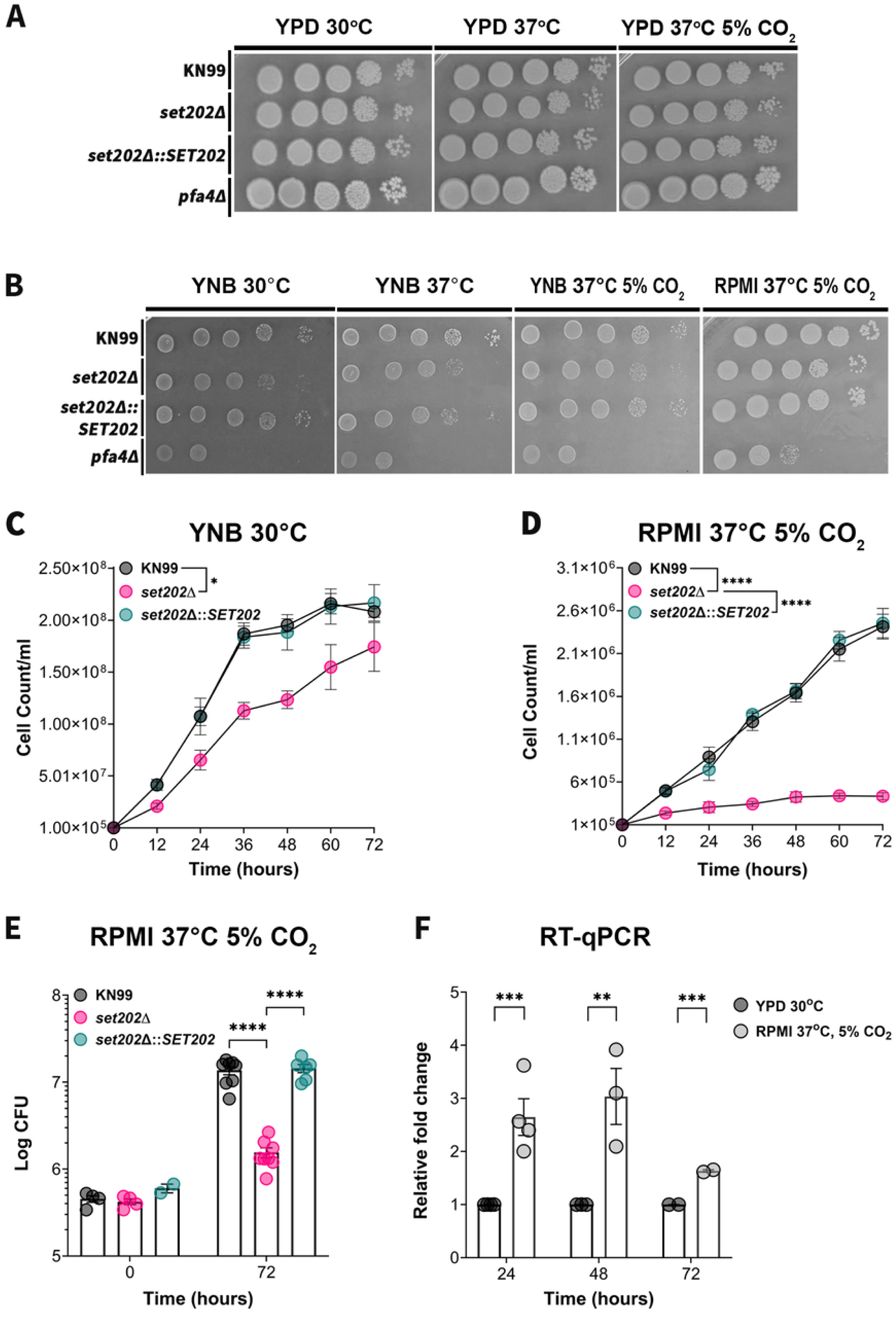
*SET202* is essential for growth in host-like conditions. (A and B) Serial dilutions of the indicated strains were spotted on YPD agar incubated at 30^°^C, 37^°^C, or 37^°^C and 5% CO_2_ (A) or on YNB agar incubated at 30^°^C, 37^°^C, or 37^°^C and 5% CO_2,_ and RPMI agar incubated at 37^°^C with 5% CO_2_ (B) and incubated for 72 hours. (C and D) The indicated strains were grown on liquid YPD media at 30^°^C (C) or RPMI media at 37^°^C and 5% CO_2_ (D) over 72 hours, and cell counts were quantified over time. (E) Samples from (D) taken at 0 and 72 hours were plated to quantify colony forming units (CFUs). (F) *SET202* relative expression in RPMI liquid media incubated at 37^°^C and 5% CO_2_ normalized to expression in YPD liquid media incubated at 30^°^C at 24,48 and 72 hours. Significance was determined by two-way ANOVA (C – E) and by T-test (F). *, P<0.05; **, P<0.01; ***, P<0.001; ****, P<0.0001.

**Fig 5.**
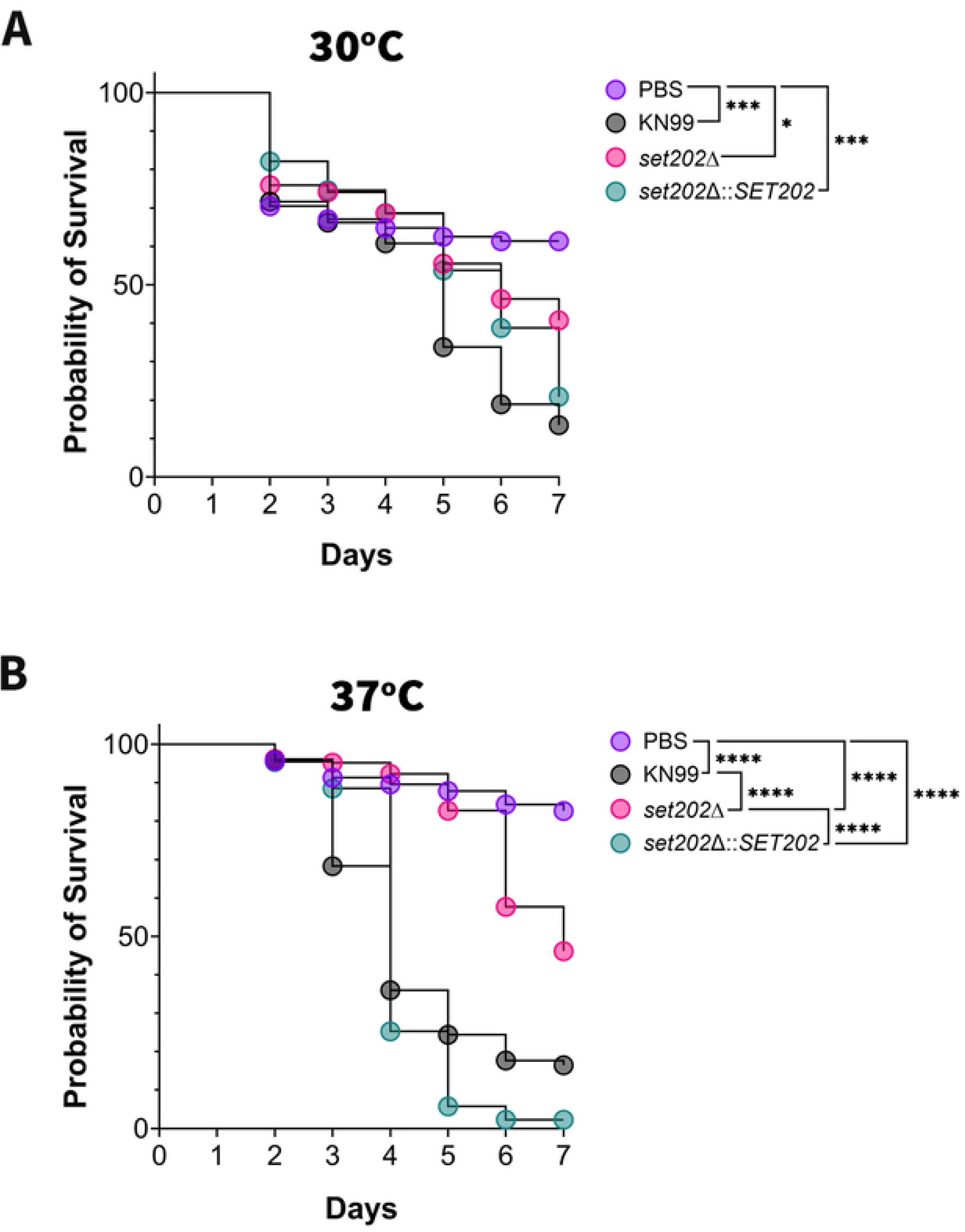
*SET202 is* essential for host-temperature-dependent fungal growth and disease progression. *Galleria mellonella* survival curves for infection with the indicated strains when incubated at 30°C (A) or 37^°^C (B) for 7days (N = 54 to 166 per group). Significance was determined by the Mantel-Cox test. *, P<0.05; ***, P<0.001; ****, P<0.0001.

If *SET202* function is needed to support normal growth under host-like conditions, we reasoned that *SET202* expression might be critical under those conditions. To assess its expression, we performed RT-PCR to measure the expression levels of *SET202* under both permissive and restrictive conditions (**Fig. 4F**). Consistently, we found a ∼3-fold increase in the expression of *SET202* when the fungi were grown in RPMI liquid media at 37 °C and 5% CO_2_ compared to YPD media at 30 °C, at 24 and 48 hours. These levels decreased slightly to ∼1.8-fold at 72hours. Together, these results suggest that Set202 plays a crucial role in the pathogenesis of *C. neoformans* by allowing the fungus to adapt and grow in the harsh host conditions that enable the establishment of infection.

### *SET202* mediates disease progression in a host-temperature-dependent manner

Thermotolerance is one of the most important fungal host adaptation mechanisms. Our findings suggest *SET202* plays a key role in fungal survival in host conditions, including higher temperatures (**Fig. 3 and 4**). To validate these observations, we repeated the infection assay in the *Galleria mellonella* invertebrate model, whose temperature we can manipulate. This model mirrors the mouse model; the fungus requires the same virulence factors to cause disease and exhibits similar immune cell-microbe interactions (28, 29). We injected the larvae with 10^5^ fungal cells and incubated them at either 30°C or 37°C over 7 days. In agreement with our murine survival data, when incubated at 37°C, *set202Δ*-infected worms showed significantly prolonged median survival of 6 days compared to 4 days for the WT and complemented-infected controls. Conversely, while there was no change in median survival at 30°C for the *set202*Δ-infected worms (still 6 days), both the WT and complemented-infected controls had their median survival increased to 6 days, as has been previously reported by other groups (29). Thus, these results show that *SET202* is required for fungal thermotolerance during infection, which is indispensable for disease causation in the mammalian host.

### *SET202* function mediates virulence factor biosynthesis in host-like conditions

In addition to thermotolerance, *C. neoformans* has several virulence factors that enable it to evade the host immune response and cause disease. Some of the major fungal virulence factors are the polysaccharide capsule, melanin, and urease. The polysaccharide capsule, composed of glucuronoxylomannan (GXM) and glucuronoxylomannogalactan (GXMGal), acts as a shield, masking fungal pathogen-associated molecular patterns (PAMPs), thereby preventing recognition, phagocytosis, and antigen presentation by immune cells (30, 31). Within the phagolysosome, immune cells generate reactive oxygen and nitrogen species (ROS/RNS) to damage the pathogen. *C. neoformans* counters this by synthesizing melanin from phenolic compounds like L-DOPA, which is abundant in brain tissue (32, 33). Urease is a fungal-secreted enzyme that acts on urea to produce carbon dioxide and ammonia, which helps the fungi neutralize the acidic phagolysosome and disseminate to the brain (34). Because thermotolerance alone does not support disease, we assessed the impact of *SET202* deletion on the production of these three major virulence factors in host conditions.

Previously, *set202*Δ mutants in the H99 background were shown to have defective melanin production at 30°C (23). While we have also found this to be true (**Fig. 6A and B**), in the KN99 background, this defect was only observed at the host temperature of 37°C. We also found that the *set202*Δ mutants in the H99 background produced no melanin at 37°C, just like a phenotypical *lac1Δ* mutant, and grew poorly compared to the *set202*Δ KN99 mutants at 37°C (**Fig. 6A and S3D**). Similarly, we found that *set202*Δ KN99 mutants produced less urease visually at 37°C compared with WT and complemented controls on urea agar (**Fig. 6C)**. As expected, there was no visual difference in induced urea breakdown at 30°C between the mutants and controls. However, this defect was not evident in urea broth at either 30°C or 37°C (**Fig. 6D**).

**Fig 6.**
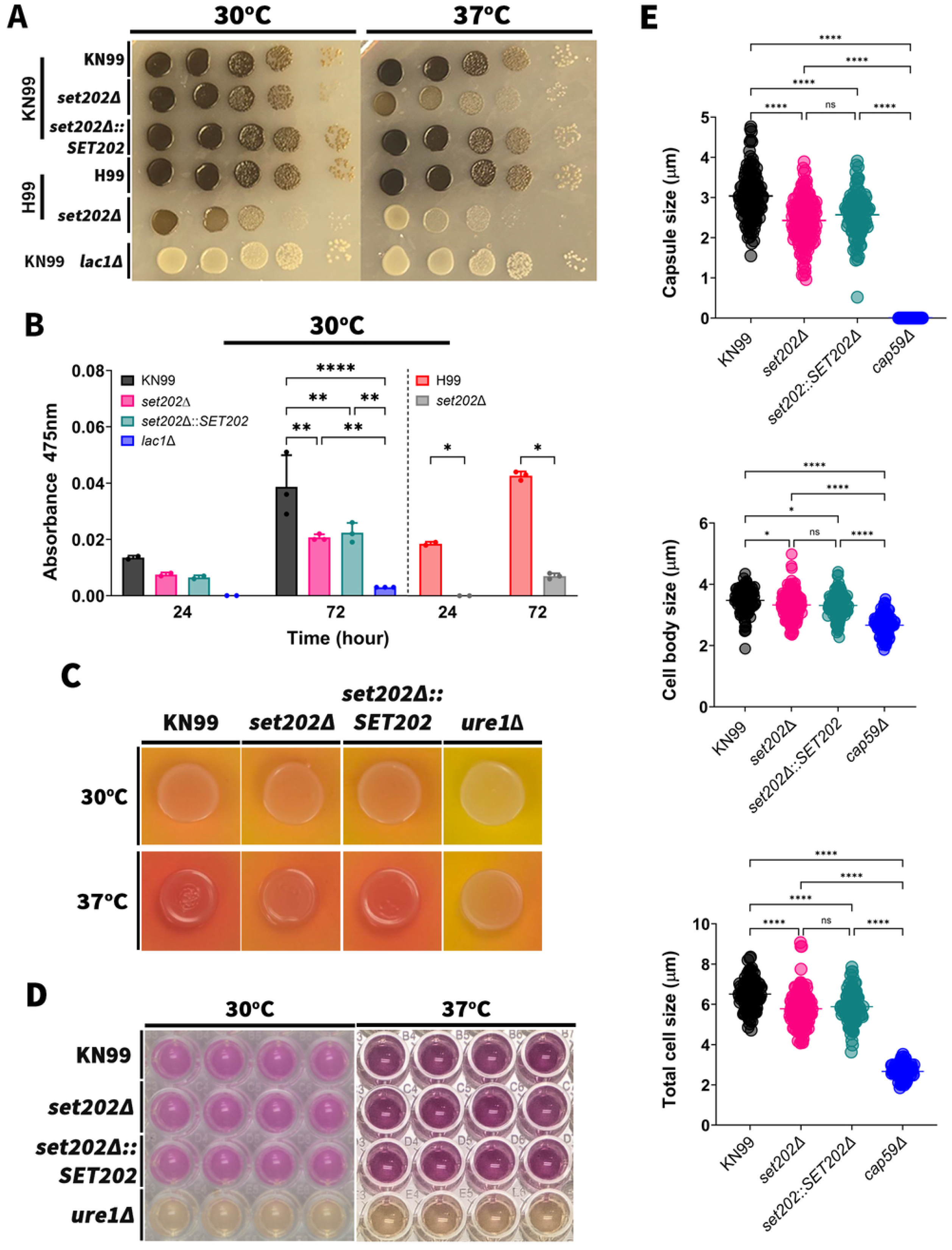
Set202 protein regulates virulence factors under host-like conditions. (A) Melanin induction assay on L-DOPA agar plates comparing the production of melanin (black pigment) among the indicated strains incubated at 30°C and 37°C for 72 hours. (B) Quantification of the melanin shed into the supernatant when growing the indicated strains in L-DOPA liquid media at 30°C for 24 or 72 hours. (C) Qualitative urease test by growing the indicated strains in urea agar for 96 hours at 30°C or 37°C. (D) Quantitative urease test by growing the indicated strains in urea liquid broth for 24 hours at 30°C or 8 hours at 37°C. In both C and D *ure1*Δ is a control strain lacking urease. (E) The indicated strains were grown in capsule-inducing media (DMEM incubated at 37°C and 5% CO_2_) over 24 hours, and the size of their capsule (top), cell body (middle), and total cell sizes (bottom) was determined. Significance was determined using one-way ANOVA with multiple comparisons. *, P<0.05; **, P<0.01; ***, P<0.01; ****, P<0.0001.

Moreover, we found that *set202*Δ mutants were hypo-capsular compared to the controls when fungi were grown in DMEM, incubated at 37°C and 5% CO_2_ for 24 hours (**Fig. 6E).** Similar findings were also seen in RPMI (**Fig. S2)**. Given the slow growth rate of the *set202*Δ mutants under host-like conditions (**Fig. 4)**, we repeated the capsule induction assay for 48 hours. Still, we found that *set202*Δ mutants were hypo-capsular despite the additional time, with no correlation between defective capsule production and cell body size (**Fig. 6D and S2)**.

Collectively, these results suggest that Set202 plays a crucial role in fungal adaptation to the host environment by supporting thermotolerance-dependent expression of major fungal virulence factors, enabling fungal survival and disease establishment.

### *SET202* does not mediate masking of fungal PAMPs under host-like conditions

The capsule plays a pivotal role in fungal virulence by masking fungal PAMPs, which promote immune evasion. Our current findings that *set202*Δ mutants are hypo-capsular (**Fig. 6**) are consistent with our prior observation that *set202*Δ mutants were easily phagocytosed compared to the WT (24). Furthermore, we find that *set202*Δ mutants are also defective in intracellular survival (**Fig. 2**). These defects may also be due to impaired masking of cell wall surface PAMPs, which would be expected with a diminished or defective capsule. Thus, we evaluated cell wall structure and surface expression of the commonly known fungal surface PAMPs beta-glucan and chitin (35).

When we tested for cell wall defects using 1 mg/ml of Congo red in YPD agar at 30°C, there were no differences in growth among the strains. However, at 37°C, we found some minor defects for the *set202*Δ mutants (**Fig. 7A**). We also found no defects at either 30°C or 37°C on YPD agar supplemented with 0.5mg/ml caffeine, an inducer of cell wall stress signaling (**Fig. 7B**). Next, we assessed surface exposure of beta-glucan and chitin when grown under host-like conditions using an antibody to beta-glucan and the dye calcofluor white. Surprisingly, despite the hypo-capsular state of the *set202*Δ mutants, we found no differences in the levels of exposure of these cell wall PAMPs (**Fig. 7C and D**).

**Fig 7.**
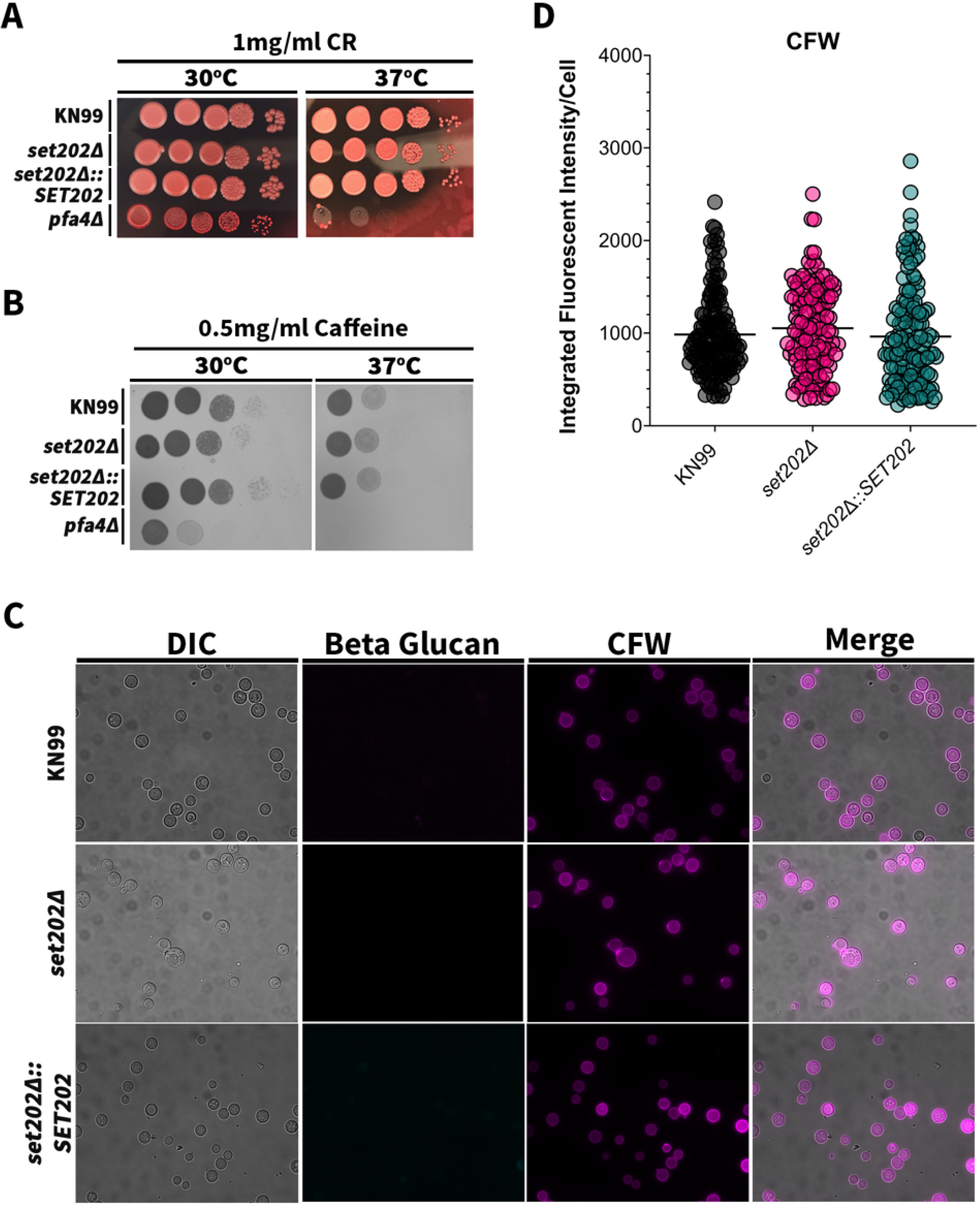
*SET202* is not involved in masking the fungal PAMPs, chitin and beta-glucan. (A and B) Serial dilutions of the indicated strains were spotted on YPD plates containing 1 mg/ml Congo Red (A) or 0.5 mg/ml caffeine (B) and incubated at 30°C and 37°C to compare growth among them. (C) Representative images of the indicated fungal cells stained for beta-glucans using an antibody or for chitin using calcofluor white (CFW) to compare the cell surface exposure of these antigens. (D) Quantification of the CFW integrated fluorescence intensity per fungal cell as a proxy of chitin cell surface exposure among indicated strains.

### *SET202* plays a role in calcium homeostasis

SET-domain HKMTs have been shown to regulate pathways that contribute to pathogenesis such as osmotic stress, cell wall integrity, and drug resistance (36, 37). In *C. neoformans*, calcium homeostasis is critical for fungal growth at mammalian host temperature (38); thus, we investigated whether *SET202* plays a role in the homeostasis of common salts, including calcium. We performed serial spot dilution assays on YPD agar plates supplemented with 1 M of various salts, incubated at either 30°C or 37°C. We found no difference in growth among the *set202*Δ mutants, WT, or complemented strains on NaCl, KCl, and MgCl_2_ (**Fig. 8A-C**).

**Fig 8.**
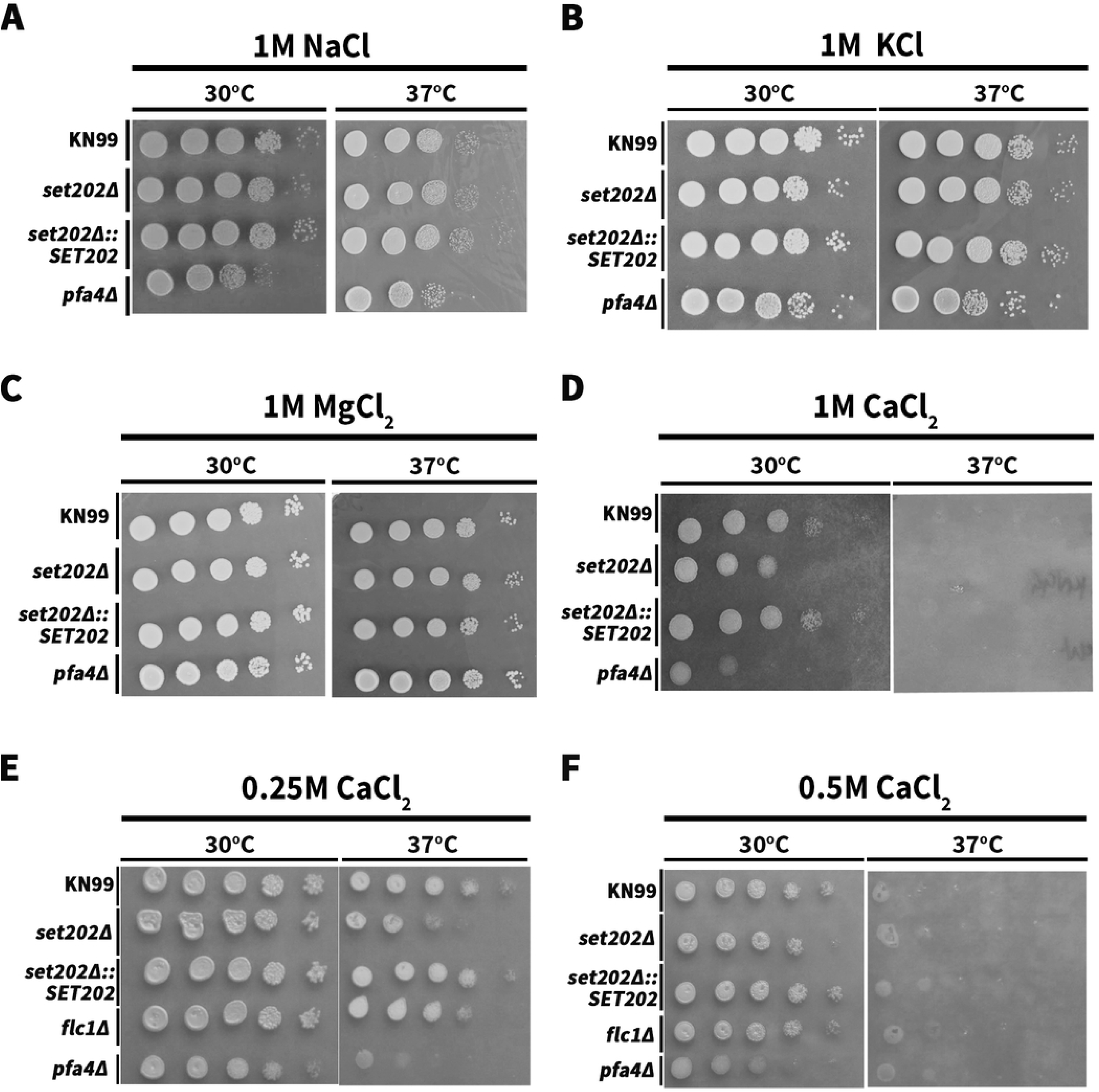
Set202 is involved in fungal calcium homeostasis. Serial dilutions of the indicated strains were spotted on YPD plates containing with 1 M NaCl (A), 1 M KCl (B), 1 M MgCl_2_ (C), 1 M CaCl_2_ (D), 0.25 M CaCl_2_ (E), and 0.5 M CaCl_2_ (F), and incubated at 30°C and 37 ^°^C for 72 hours.

Interestingly, on the 1 M CaCl_2_ agar plates at 30°C, the *set202*Δ mutants showed significantly less growth than the WT and complemented controls (**Fig. 8D**). Also, the *set202*Δ mutants were slightly more susceptible to calcium than the *flc1Δ* mutants, which have previously been described as having defective calcium homeostasis (**Fig. S3A**) (39). Since none of the mutants grew on the 1 M CaCl_2_ agar plates incubated at 37°C (**Fig. 8D**), we tested lower concentrations (**Fig. 8E and F**). We found no overt differences in growth among *set202*Δ mutants, WT, or complemented strains on 0.25 M CaCl_2_ incubated at 30°C. However, the *set202*Δ mutants showed defective growth at the same calcium concentration when incubated at 37°C. This growth defect is independent of temperature, as they grow fine in YPD media, with or without other salts, at 37°C (**Figs. 4A and 8A – C**). Similar to the 1 M CaCl_2_ phenotype, *set202*Δ mutants grew poorly on 0.5 M CaCl_2_ plates incubated at 30°C, and at 37°C, all the strains showed very little growth.

Collectively, these results show that beyond adaptation to host conditions, *SET202* also plays a key role in fungal calcium homeostasis in a temperature- and dose-dependent manner, which could contribute to the impaired survival of the *set202*Δ mutants in the host.

#### Phylogeny, Protein sequence, and predicted structure

SET domain proteins are conserved across the fungal kingdom, where they play important roles in epigenetic modifications (40). Among pathogenic fungi, they have been associated with survival during nutrient stress, antifungal resistance, and disease dissemination (41, 42). Although Set202 has been proposed to be a putative histone lysine methyltransferase, no group has tested this (23). This potential function was based on sequence homology; thus, we decided to investigate this further.

An NCBI BLAST search was used to generate a phylogenetic tree of fungal histone 3 lysine 36 (H3-K36) SET-domain-related genes (**Fig. 9A**). Using a threshold of 30% sequence identity to the *SET202* gene in KN99 (highlighted green), the tree includes many common pathogenic fungi that are part of the WHO’s fungal priority pathogen list (43), as well as the model yeast *Saccharomyces cerevisiae* (44), highlighted in orange.

**Fig 9.**
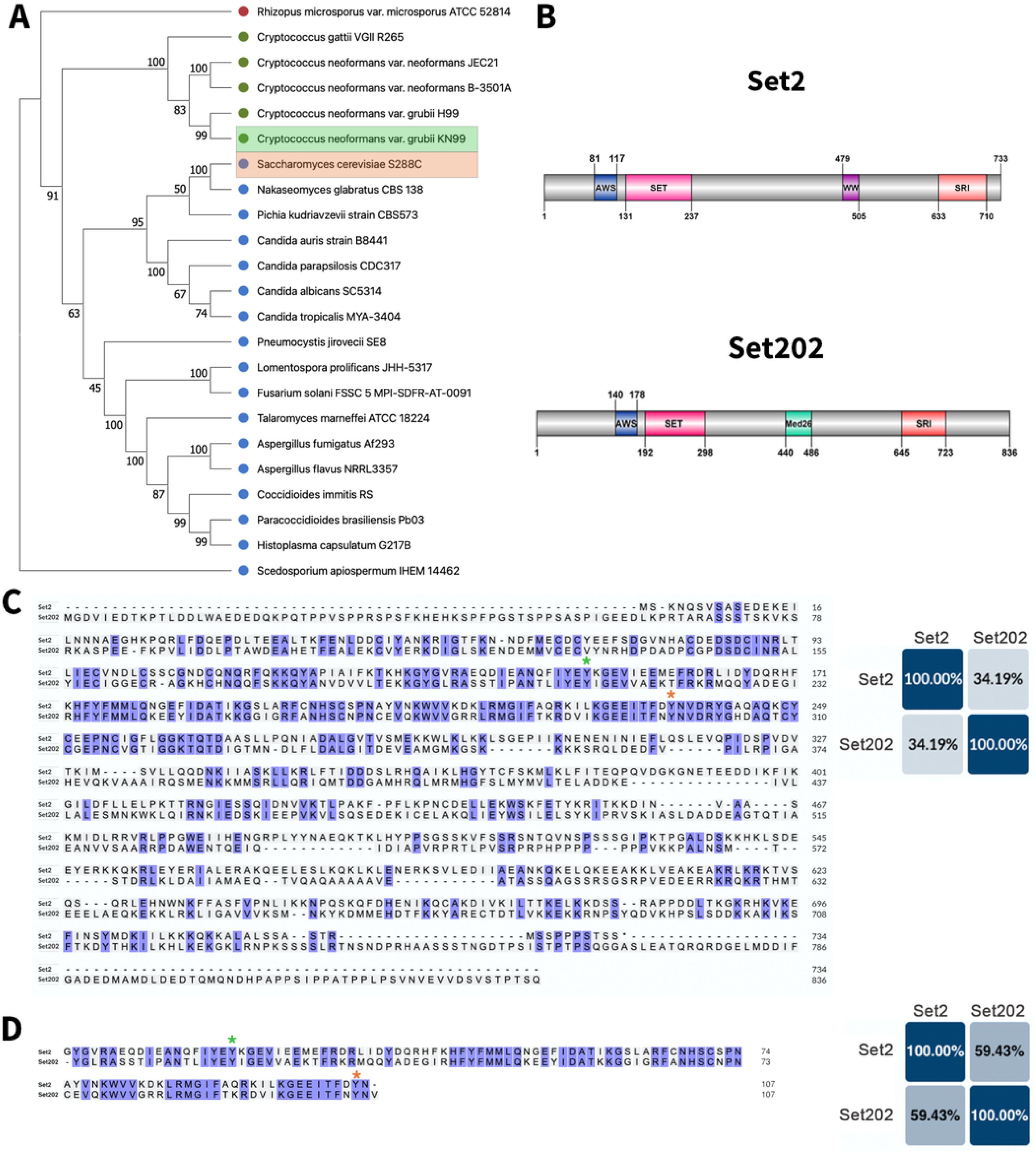
*SET202* Phylogeny and predicted protein domains. (A) Phylogenetic tree of *SET202* (highlighted in green) homologs in common pathogenic fungi and *S. cerevisiae* (*SET2* highlighted in orange). The gene sequences used are from the specified genomes following FungiDB nomenclature. (B) Predicted protein domains of Set2 and Set202 according to InterPro and illustrated using DOG 2.0 (59). (C) Uniprot sequence alignment and percent identity of Set2 and Set202. (D) Uniprot sequence alignment and percent identity of the predicted SET-domain catalytic site of Set2 and Set202. Highlighted in C and D with asterisks are the catalytic sites Y149 (green) and Y236 (orange) of Set2.

An InterPro comparison of Set202 to its *S. cerevisiae* ortholog, Set2, shows similar architecture and protein domains (**Fig. 9B).** Both contain AWS (Associated with SET) domains, catalytic SET domains, and SRI (Set2 Rpb1 Interacting) domains. Full-length sequence alignment of both proteins on Uniprot showed 34.19% identity, as previously predicted by our BLAST search **(Fig. 9C)**. If we restrict the alignment to the catalytic SET domain, which is closely related in different organisms (40, 45), the sequence identity increased to 59.43% (**Fig. 9D**). Set2 is a known H3-K36 methyltransferase (H3K36MT) responsible for three methylation activities on the histone 3 tail: mono-methylation (H3K36Me1), di-methylation (H3K36Me2), and tri-methylation (H3K36Me3)(42). At its SET domain catalytic site, mutation of Y149 to phenylalanine significantly decreased Set2-mediated H3K36Me1 activity and completely abolished its H3K36Me2 function. A similar mutation at Y239 resulted in the complete loss of Set2’s H3K36Me3 (46). Notably, our sequence alignment shows that Y149 and Y239 correspond to Y210 and Y297, respectively, in Set202, supporting the evolutionary conservation of the SET-domain-containing proteins and the possibility that Set202 might exhibit the same activity.

### Set202 is a bona fide histone 3 lysine 36 methyltransferase

To determine whether Set202 also mediates H3K36Me1, H3K36Me2, and H3K36Me3 posttranslational modifications (PTMs), we probed total protein lysates from all fungal strains with antibodies specific for histone 3 and for these modifications, with beta-actin as normalization and loading control. Interestingly, we found a minor decrease in the overall expression level of histone 3 in *set202*Δ mutants compared to the WT and complemented controls (**Fig. 10A**). Excitingly, we found significantly reduced levels of the three methylation activities in *set202*Δ mutants (**Fig. 10 B-D**), supporting the notion that Set202 is an H3K36MT.

**Fig 10.**
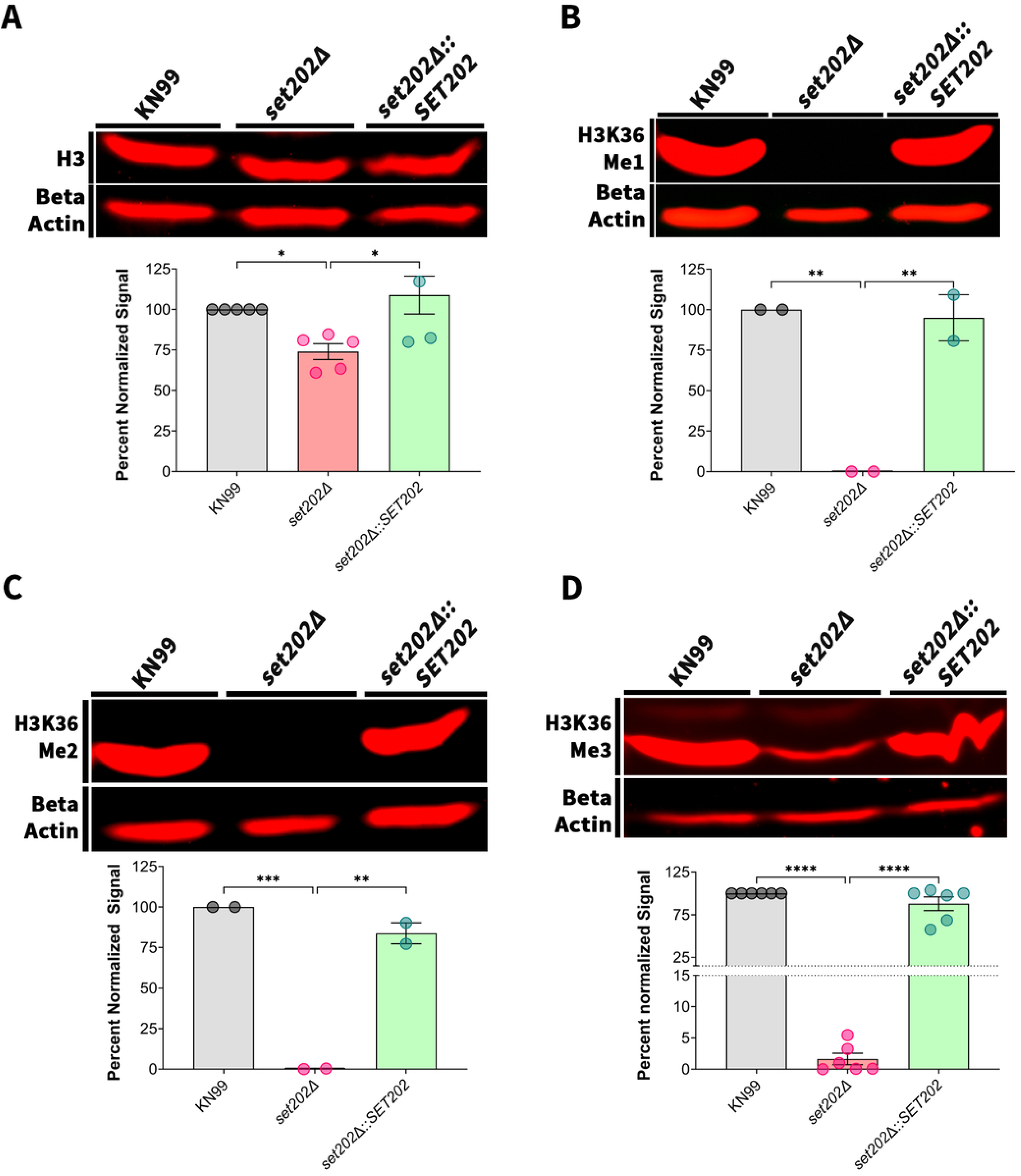
Set202 protein is a histone-3 lysine-36 (H3K36) methyltransferase. (A – D) Immunoblotting of whole-cell protein extracts from the indicated strains showing the protein levels of A) total histone-3, B) histone-3 lysine-36 mono-methylated, C) histone-3 lysine-36 di-methylated, and D) histone-3 lysine-36 tri-methylated. Beta-actin was used as a loading control. Quantification of the bands was done by normalizing to beta-actin. Significance was determined using one-way ANOVA. *, P<0.05; **, P<0.01; ***, P<0.01; and ****, P<0.0001.

Collectively, our data suggest that Set202 functions as an H3K36MT that epigenetically regulates gene expression, enabling *C. neoformans* to rapidly adapt to the host internal milieu and cause disease. In line with these conclusions, we propose a dual mode of action for Set202, regulating gene expression in response to environmental or host stressors (**Fig. 11**).

**Fig 11.**
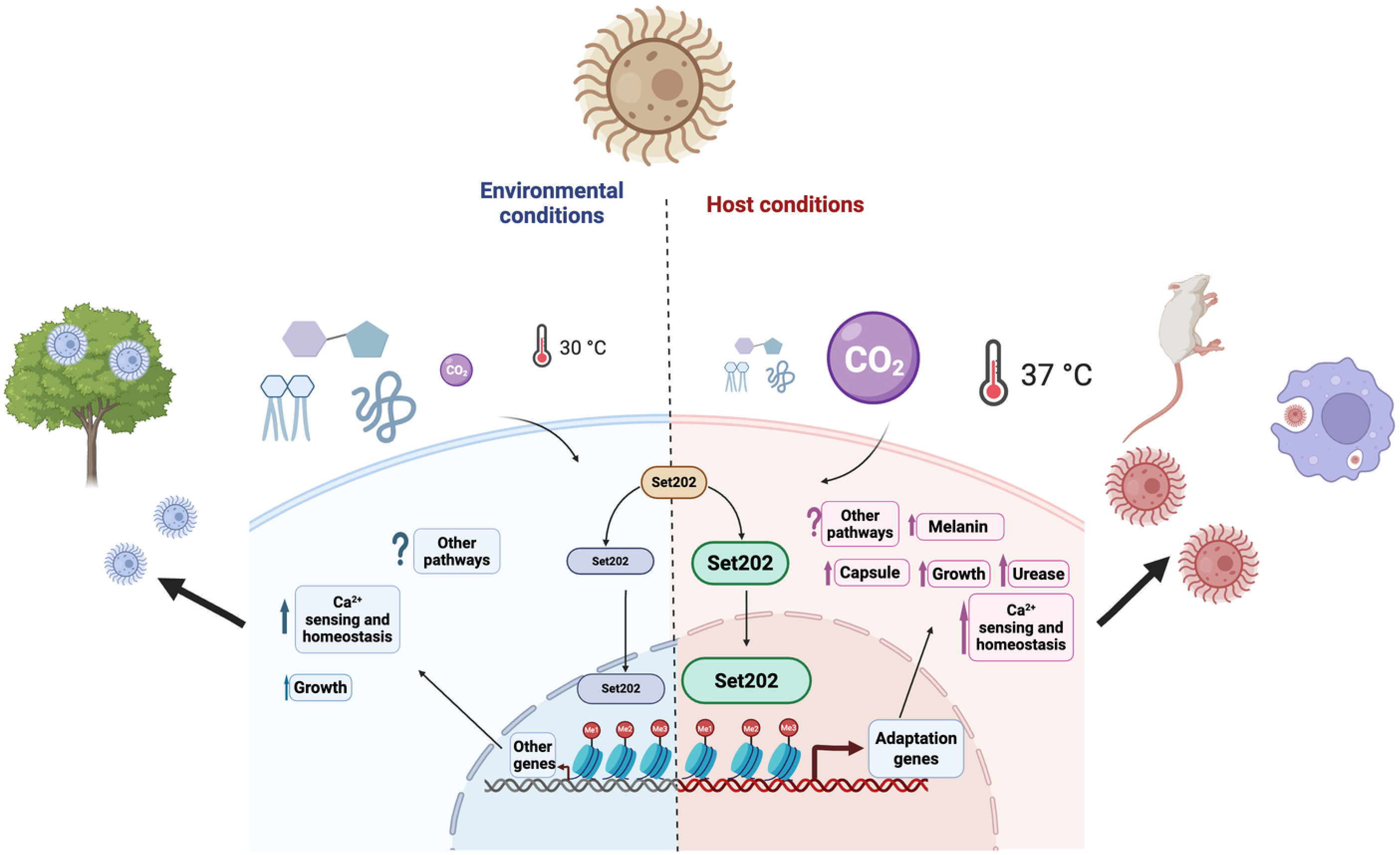
Model of function of *Set202* and role in fungal host adaptation mechanisms. Model summarizing the function of *SET202* in *C. neoformans* host-pathogen interactions. Under environmental conditions (low carbon dioxide concentrations, nutrient excess, and ambient temperature), Set202 mainly mediates fungal calcium sensing and homeostasis. However, under host conditions (high carbon dioxide concentrations, limited nutrient excess, and a temperature of 37°C), Set202 upregulates fungal adaptation and virulence genes that promotes growth, impair immune responses, and mediate faster disease progression. Figure created in BioRender (https://biorender.com/clqnrnm).

## DISCUSSION

Mechanisms of fungal adaptation to mammalian host conditions are central to the pathogenesis of environmental pathogens such as *C. neoformans*. These adaptive responses rely on rapid molecular changes at the genome level, which involve histone PTMs such as histone methylation. SET-domain histone methyltransferases have been implicated in fungal virulence processes, such as drug resistance, phagocytosis, and thermotolerance. Here, we identify Set202 as an H3K36 methyltransferase that regulates the adaptation of *C. neoformans* to host conditions.

*SET202* was first identified as a gene required for growth in murine lungs and named after its homolog in the model yeast *S. cerevisiae*, Set2. However, despite being identified in other studies related to pathogenesis, including in this study as a regulator of phagosomal permeabilization, the cellular and molecular function of Set202 remained unknown. Comparative analysis revealed that Set202 shares conserved AWS, SET, and SRI domains with Set2, a canonical H3K36 methyltransferase (18), with especially high sequence identity at the predicted catalytic sites within the SET domain. In line with these structural similarities, our biochemical assays demonstrate that Set202 functions as a true H3K36 methyltransferase in *C. neoformans*, highlighting that H3K36 methylation has a conserved role in regulating transcriptional machinery during stress across divergent fungal lineages.

In *S. cerevisiae,* Set2-mediated H3K36 methylation is crucial for stress adaptation, including nutrient deprivation, genotoxic and DNA damage, and ageing (18). For pathogenic fungi like *Cryptococcus*, host conditions such as elevated temperature, limited nutrients, and increased carbon dioxide tension represent significant sources of cellular stress; thus, it is reasonable to hypothesize that histone methylation might be implicated in stress adaptation in this fungus too. Indeed, we find that in the absence of Set202-mediated H3K36 methylation, the cells displayed no or mild growth defects under homeostatic conditions (i.e., YPD or YNB at 30°C), but they showed pronounced growth defects under host-like conditions (RPMI medium at 37°C and 5% CO_2_). Furthermore, we found that, under these conditions, the *set202*Δ mutants are defective in several of the main virulence factors of *C. neoformans*, which are also induced upon exposure to host conditions.

Two of the most important *C. neoformans* virulence factors are melanin, a reactive oxygen species scavenger, and the immunomodulatory polysaccharide capsule, both of which are tightly regulated by the cellular environment (47, 48). Interestingly, other histone PTMs have been shown to contribute to regulation of these factors. The Set302 protein, a histone deacetylase complex, is required for capsule growth under host-like conditions and for melanin production at 39°C, the avian reservoir temperature (49); and Set1, part of the COMPASS complex that also carries out histone PTMs, is needed for full melanization at 30°C (50). Similarly, Set202 has a role in these processes but only under host-like conditions, a finding that explains a prior report. In the clinical H99 strain, Set202 is needed for melanin production starting at 30°C, a defect that was more pronounced at the mammalian host temperature of 37°C (23). In contrast, in the lab-adapted KN99 strain, Set202’s role in melanin biosynthesis is only evident at 37°C. These observations suggest that similar SET-domain proteins within the same species can function in a strain-specific, temperature-dependent manner, attesting to the evolutionary fine-tuning of the function of these proteins in response to diverse host stressors.

These *in vitro* phenotypic defects clearly alter host-pathogen interactions important for disease. Set202 contributes to phagosomal permeabilization and delayed recruitment of the v-ATPase acidification complex, thus promoting intracellular survival. In its absence, *set202*Δ mutants are rapidly killed by macrophages. This is consistent with our prior report of increased phagocytosis of *set202*Δ mutants by THP-1 macrophages (24), given their smaller capsule. Thus, it is not surprising that *SET202* deletion results in markedly attenuated disease progression in murine and waxworm infection models, similar to what has been reported for SET-domain proteins in other pathogenic fungi (21, 22, 49). For example, in *Candida albicans*, Set1-mediated H3K4 methylation accelerates disease progression by dysregulating mitochondrial ROS metabolism and increasing survival in phagocytes (20, 22). Interestingly, using the waxworm model of infection allowed us to test the effect of temperature on disease, finding that Set202-mediated infection and death in the waxworm model are temperature-dependent, similar to other fungal virulence genes (29, 51, 52). Another notable observation was the rapid, albeit temporary, seeding of the brain by the *set202*Δ mutants in our murine model. The smaller cell sizes of the *set202*Δ mutants may explain the day 7 brain dissemination, as a smaller morphotype of *C. neoformans* called seed cells has been reported to exhibit increased and faster dissemination from lungs (5). However, the fact that *set202*Δ mutants are easily internalized could result in direct fungal transit to the brain via the cribriform plate during the intranasal infection, as suggested recently in a study (53). However, once in the brain, these cells are rapidly cleared, and it is not until 4 weeks later that we see reseeding of the brain, which then keeps increasing over time until the time of death. This phenomenon can be explained by either (i) selection of a subset of better-adapted *set202*Δ mutants from the stressors in the pulmonary environment, or that other fungal-adaptation pathways were upregulated later by the fungi, in lieu of *SET202* deletion, or (ii) the increased burden in the lungs results in an increased dissemination rate which overwhelms the brain defenses. Given that calcium signaling plays a key role in stress adaptation in *C. neoformans* (54), and that *set202*Δ mutants were sensitive to lower and higher concentrations of calcium at 37°C and 30°C, respectively, we cannot rule out that the eventual lethal progression of the disease is due to delayed activation of alternative compensatory stress-response pathways in the absence of *SET202*.

In conclusion, we have uncovered a new mechanism by which *C. neoformans* adapts to host conditions, via the H3K36 methylation activities of the Set202 protein. While our results provide a cellular and molecular function to Set202, it will be interesting to find downstream affected genes, link them to our current findings, and develop a signaling pathway that correlates these. Our work supports the role of Set202 in mediating chromatin remodeling and epigenetically activating relevant transcriptional adaptation genes, required for fungal survival under the complex stressors found in the host. This knowledge will outline one of the mechanistic pathways by which *C. neoformans* adapts to host conditions and lay the groundwork for understanding fungal host-pathogen interactions, and possibly new therapeutic targets.

## MATERIALS & METHODS

### Fungal Strains

*C. neoformans* strains (either in the KN99α or H99 background) were obtained from the Madhani collection (23), and the *S. cerevisiae* strain BY4741 served as a control. All strains came from YPD-glycerol stocks at -80°C, streaked onto YPD agar, stored at 4°C, and replaced every 6 months from the -80°C stocks.

### Complement construction

*SET202* complemented strains were constructed using a modified Transient CRISPR-Cas9 Coupled with Electroporation (TRACE) method, as described (55). Briefly, the *SET202* coding sequence was PCR-amplified from genomic DNA using primers with NotI and KpnI restriction sites, cloned into pQE-NEO_E2 plasmid, and transformed into *E. coli* DH5α. This plasmid was used to obtain a repair construct containing *SET202* and the neomycin resistance cassette by PCR. This construct was targeted to the Safe Haven 2 locus (SH2) by electroporation together with Cas9 and SH2 sgRNA. Transformants were selected on YPD with 100ug/ml neomycin, verified by PCR, and further assessed for complementation by restoration of melanin production at 37°C. Primers, restriction enzymes, and plasmids used are listed in **Table S2**. For more details on these procedures, see Supplementary Text S1.

### Growth assays and stress plates

Overnight YPD cultures (30°C) were washed and counted with a Bio-Rad TC20^TM^ Automated Cell Counter. Equal cell counts were inoculated into the indicated liquid media for growth curve assays. For spot assays, serial dilutions were prepared from normalized cultures, and 5 µL of each dilution was spotted for each strain on the indicated agar plates. Plates were incubated as specified on figure legends. Additionally, cell counts were quantified at specified time points, and colony-forming units (CFUs) were determined at the initial and final time points to confirm viability during growth in liquid RPMI incubated at 37°C with 5% CO_2_.

### Capsule induction and cell wall stains

Equal fungal counts were inoculated into DMEM or RPMI for cell wall staining and capsule induction, or 10% heat-inactivated fetal bovine serum for titan cell induction. Cultures were incubated at 37°C with 5% CO_2_ for 24 - 72 hours depending on the assay. Titan cell and capsule induction were assessed using Indian ink; cell wall chitin was visualized with calcofluor white; and beta-1,3-glucan exposure was assessed with anti-beta-glucan antibody. Imaging was performed using a 100x oil-immersion objective on a Zeiss Axio Observer 7 microscope. Quantification of capsule size, cell wall staining, and titan cell sizes was done using FIJI or a Cell Profiler pipeline. Automated quantification of titan cell size was also obtained using the FIT Flow CAM VS-1-C B3 instrument from the Center for Environmental Science and Technology (CEST) at the University of Notre Dame. For more details on these procedures, see Supplementary Text S1.

### Cell culture, immunofluorescence and phagocytosis/survival assays

THP-1 cells were maintained in RPMI 1640-based media as described (24). Cells were passaged at least twice weekly and were not used beyond passage 15. For differentiation into macrophages, PMA was used as described (56, 57). THP-1 cells were seeded at 1.5 × 10^5^ cells/mL in a 24-well plate containing 12-mm round glass coverslips for immunofluorescence or without coverslips for killing assays. Fungal strains were stained with Lucifer Yellow, opsonized with normal human serum, and added to the THP-1 cells at a multiplicity of infection (MOI) of 1:1. At the indicated time points, coverslips were washed twice with PBS and fixed with formaldehyde for 10min, then processed immediately or stored at 4°C until staining. Preparation for microscopy was done as previously described (24). Primary antibodies were diluted in 1% goat serum or BSA and applied as follows: anti-galectin-3 antibody (1:100; overnight), human anti-LAMP-1 (1:100; overnight) and anti-vATPase (1:500; 1 hour). Imaging was performed using a 100x oil-immersion objective on a Zeiss Axio Observer 7 microscope and antibody-positive phagosomes was determined using FIJI and plotted in GraphPad Prism v10. For more details on these procedures, see Supplementary Text S1.

### Animal infection models

Healthy larvae weighing >190mg were acclimated at room temperature for 24 hours before infection by injecting 5 µL of sterile PBS or 1 × 10^5^ stationary-phase fungal cells in the posterior proleg as described (58). Larvae were incubated at either 30°C or 37°C and monitored daily for survival over 7 days. For the murine model, we used 5-to 6-week-old female A/J mice as described (56, 57). Mice were infected intranasally with 5 × 10^5^ log-phase fungal cells in 40 µL of sterile DPBS or received 40 µL DPBS alone. Animals were monitored daily and euthanized upon losing >20% of peak weight. Brain, lungs, and spleen were harvested, homogenized in 5 mL DPBS, and plated on YPD with 1 µg/mL ampicillin (30°C, 48 h) to quantify CFUs. All murine procedures were approved by the University of Notre Dame Institutional Animal Care and Use Committee (protocol #23-05-7918). For more details on these procedures, see Supplementary Text S1.

### RNA preparation, quantitative real-time PCR, and Western blotting

For RNA isolation, cells were grown in YPD at 30°C or RPMI at 37°C with 5% CO_2,_ harvested, resuspended in TRIzol reagent, and disrupted by rigorous bead-beating using a Mini-beadbeater-16. RNA was extracted via chloroform/isopropanol/ethanol treatment, purified using Zymo RNA Clean & Concentrator-25, and reverse-transcribed with SuperScript III. Real-time PCR was performed using SsoAdvanced Universal SYBR Green Supermix on a Bio-Rad Real-Time PCR system and gene-specific primers (see **Table S2**) loaded into a 96-well plate. Data were analyzed using Bio-Rad PrimePCR Analysis software and GraphPad Prism v10.

For protein isolation and immunoblotting, YPD-grown log phase cells were harvested, resuspended in ice-cold lysis buffer with protease inhibitors, and disrupted by bead-beating as above. Lysates were incubated at 4°C with 1% Triton X, clarified by high-speed centrifugation, and total protein quantified via BCA assay. Protein samples were stored at -20°C or -80°C for long-term storage. Equal amounts of protein were resolved on SDS-PAGE gel and transferred to PVDF membranes. Membranes were blocked in 5% milk/TBS for 1 hour, incubated overnight at 4°C with primary antibodies in 3% milk/TBS-T (anti-beta actin, anti-Histone-3, anti-Histone-3-Lysine 36 mono-methylation, anti-Histone-3-Lysine 36 di-methylation, and anti-Histone-3-Lysine 36 tri-methylation), and after washing (TBS-T with 0.01% SDS), probed with secondary antibodies (IRDye 680RD) in 3% milk/TBS-T with 0.01% SDS. Membranes were imaged in an Odyssey CLx imaging system, band intensities quantified using Image Studio v5.2, and data plotted in GraphPad Prism v10. For more details on these procedures, see Supplementary Text S1.

## ACKNOWLEDGMENTS

We are grateful for the support provided for the murine studies by the staff from the University of Notre Dame Freimann Life Science Center (FLSC). Special thanks to the members of the Santiago-Tirado, Flores-Mireles, Wingert, Schafer, McConnell, Shrout, and Champion laboratories for their constructive feedback, sharing equipment, and sharing reagents. This work was supported in part by a University of Notre Dame Center for Environmental Science and Technology (CEST) Predoctoral Fellowship (to GA), the Berthiaume Institute for Precision Health Summer Graduate Fellowship (to GA), the ASPiRE-ND Prep program (to KCA), institutional funds (to FHS-T), and NIH grants (to FHS-T; R21AI71742 and R01AI177875).

